# Discrete Diffusion for Single-Cell Gene Expression Modeling

**DOI:** 10.64898/2026.02.19.705033

**Authors:** Sanjukta Bhattacharya, Christian Gensbigler, Shaamil Karim, Jon Lees

## Abstract

Current generative modeling of single-cell transcriptomics relies on continuous latent representations, transforming inherently discrete and sparse gene counts into continuous space. We propose Discrete Cell Models (DCM), a diffusion-based framework that learns cellular representations directly in the discrete domain. Our framework supports both unconditional and conditional generation, allowing for precise modeling of complex biological scenarios such as cell-type-specific transcriptional responses to genetic perturbations. We demonstrate that DCM scales effectively and achieves strong performance against current state-of-the-art methods, including scVI, CPA, STATE, scGPT, and scLDM. On the Dentate Gyrus benchmark, DCM achieves a 5-fold improvement in *MMD*^2^*RBF* and a nearly 2-fold improvement in *W*_2_ distance, over the leading continuous diffusion baseline (scLDM). On the conditional Replogle perturbation benchmark, DCM sets a new state of the art on *W*_2_ distance while remaining competitive on *MMD*^2^*RBF*. Together, these results establish discrete diffusion as a promising direction for foundational models of cellular biology.

## 1 Introduction

A cell can be viewed as an information-processing system that maps environmental inputs to molecular responses through an internal regulatory program Fitch (2021). While the structure of the regulatory network itself is not directly accessible, single-cell RNA sequencing provides a discrete high-dimensional but partial observation of its activity Macosko et al. (2015); Gulati et al. (2020); Zeng & Dai (2019). Crucially, this observation is discrete: we do not measure continuous expression levels, but count individual molecular events. Across many cells and conditions, these observations implicitly encode the statistical dependencies induced by the underlying regulatory structure. The objective of a virtual cell model is to learn these dependencies well enough to generate realistic cellular states and to predict how modifications to the regulatory program, such as genetic perturbations, alter the resulting transcriptome Dixit et al. (2016); Adamson et al. (2016).

The central object of these observations is the count matrix: a sparse, integer-valued tensor recording how many mRNA molecules of each gene were captured from each cell. Yet the dominant generative models have been applied to this problem begin by embedding these integers into continuous vector spaces Lopez et al. (2018b); Gandhi et al. (2025); Bereket & Karaletsos (2023); Adduri et al. (2025). Variational autoencoders such as scVI Lopez et al. (2018a); Kingma & Welling (2013) learn low-dimensional latent representations under a negative binomial observation model, accounting for the count nature of the data at the likelihood level but restricting the latent prior to a unimodal Gaussian. Compositional Perturbation Autoencoder (CPA) Lotfollahi et al. (2023) extends this framework by disentangling cell identity from perturbation effects in latent space, enabling combinatorial prediction of unseen conditions. Transformer-based foundation models like scGPT Cui et al. (2024) scale to tens of millions of cells and produce powerful embeddings. Although operating in the discrete space, scGPT’s autoregressive generation imposes a sequential ordering on genes, which is an inductive bias that contrasts with the exchangeability of gene expression Kim et al. (2025).

More recently, diffusion-based approaches have addressed generation more directly. scDiffusion Luo et al. (2024) applies classifier-guided continuous diffusion in the latent space of a pre-trained foundation model. CFGen Palma et al. (2024) introduces flow matching Lipman et al. (2022) with compositional guidance and explicit negative binomial likelihoods, enabling generation conditioned on multiple biological attributes simultaneously. scLDM Palla et al. (2025), the current state of the art, combines a fully transformer-based VAE, with permutation-invariant encoding and equivariant decoding, with a latent diffusion model parameterized by Diffusion Transformers Peebles & Xie (2023) and trained via flow matching. scLDM demonstrated that architectural respect for the exchangeability of genes yields substantial gains in generation fidelity. Yet despite their differences in architecture and training objective, all of these recent methods share a common representational commitment: they transform raw integer counts into continuous vectors and model in the resulting real-valued space. The discrete structure of the raw measurements is recovered only after sampling, through rounding or sampling from count distributions fitted to continuous predictions.

This representational choice warrants scrutiny. In language modeling, we do not embed discrete tokens into continuous space, generate in that space, then discretize back—we model tokens directly Vaswani et al. (2017). Yet for transcript counts, the field has adopted continuous relaxation as a default Lopez et al. (2018b); Gandhi et al. (2025); Bereket & Karaletsos (2023); Adduri et al. (2025). The standard justification may be that continuous spaces are easier to model, and that the observation model (Poisson, negative binomial) accounts for discreteness at the likelihood level. We argue that this justification is insufficient, and that continuous relaxation introduces fundamental representational limitations.

First, continuous models assign probability mass to non-integer values that correspond to no possible measurement—model capacity spent on impossible states. Second, the natural metric on count space is not Euclidean: the difference between 0 and 1 transcript (presence versus absence of expression) is biologically distinct from the difference between 100 and 101 transcripts (sampling noise) Hafemeister & Satija (2019). Continuous embeddings with Euclidean metrics cannot represent this asymmetry without learning it from data; discrete models respect it by construction. Third, gene regulatory networks induce relations that depend on the functional presence of gene products, which for lowly-expressed genes is inherently stochastic and discrete—a gene with mean expression 0.5 transcripts/cell is “off” in most cells and “on” in some, a bimodal phenomenon that continuous models must learn to represent but that discrete models capture naturally Elowitz et al. (2002). Finally, there is an information-theoretic argument: when the true data-generating process is discrete, any continuous relaxation introduces a representation gap, forcing the model to learn discretization boundaries rather than structure within the discrete space Maddison et al. (2016). Furthermore, recent advances in discrete diffusion for text Lou et al. (2023); Sahoo et al. (2024) and protein sequences Tang et al. (2024); Gruver et al. (2023) have demonstrated that operating directly in discrete token space yields substantial improvements over continuous relaxations—results that suggest similar gains should be achievable for the discrete, sparse count distributions characteristic of single-cell transcriptomics.

We introduce Discrete Cell Models (DCM), a framework that applies Score Entropy Discrete Diffusion (SEDD) Lou et al. (2023); Campbell et al. (2022) directly to raw transcript counts, eliminating the continuous relaxation used by prior work. DCM treats each gene’s expression level as a discrete token and learns to reverse a corruption process over the count space via the concrete score: the ratio of data distribution across neighbouring discrete states Meng et al. (2022). The model supports multi-conditional and unconditional generation within a single end-to-end architecture, synthesizing cell states conditioned jointly on cell type, perturbation identity, and other biological attributes. We evaluate DCM on two complementary benchmarks. On the Dentate Gyrus dataset (unconditional generation)Hochgerner et al. (2018), DCM achieves a nearly 2-fold reduction in *W*_2_ distance and a 5-fold improvement in *MMD*^2^*RBF* over scLDM. On the Replogle dataset (conditional perturbation prediction), DCM achieves the best *W*_2_ distance across all baselines while maintaing a competitive *MMD*^2^*RBF*. Together, these results establish discrete diffusion as a promising direction for foundational models of cellular biology.

## 2 Discrete diffusion for gene expression generation

### Gene Expression Representation

We represent single-cell gene expression profiles as discrete sequences. Let **x** ∈*χ*^*M*^ denote a cell’s expression profile across *M* genes, where *χ* = {0, 1, …, *K*} is the discrete vocabulary of binned or raw expression counts, with *K* as the max gene expression count in the dataset. Each element *x*_*i*_ ∈ *χ* represents the expression level of gene *i* in a single cell.

**Figure 1.**
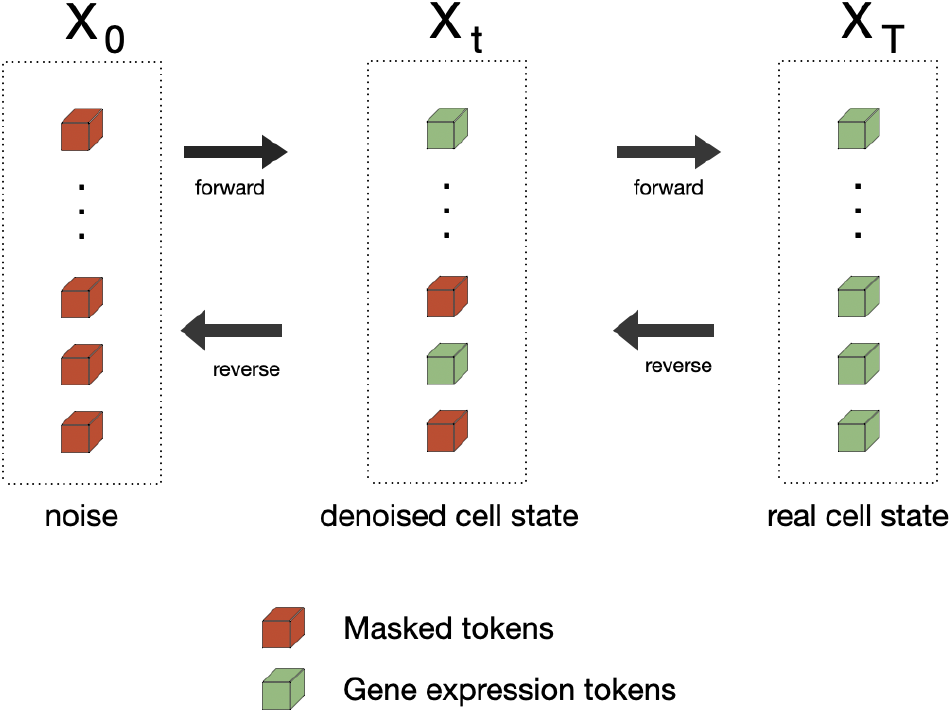
Generating single cell data with DCM

### Conditional Generation Objective

We aim to learn a generative model *p*_*θ*_(**x**|**c**) that produces gene expression profiles conditioned on cellular attributes **c** = (pert, cell type), where pert denotes the genetic perturbation (e.g., gene knockdown) and cell type specifies the cellular context (e.g., HepG2, Jurkat).

Given observed i.i.d. data 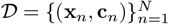, where each observation consists of a gene expression profile **x**_*n*_ and its corresponding conditions **c**_*n*_, our objective is to learn a parametric model *p*_*θ*_(**x** |**c**) with parameters *θ* that maximizes the conditional log-likelihood:

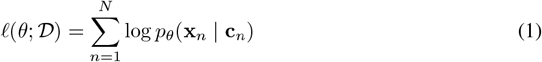

### Discrete Diffusion Framework

We use Score Entropy Discrete Diffusion (SEDD) Lou et al. (2023), which models the data distribution through a continuous-time discrete-state markov process. Rather than directly parameterizing *p*_*θ*_(**x** |**c**), SEDD learns to estimate ratios of the data distribution at different noise levels which is then used to sample the gene expressions.

### Forward Diffusion Process

We define a continuous-time markov process that progressively corrupts clean expression profiles **x**^0^ ∼ *p*_data_(**x** | **c**) through independent token-level transitions. For time *t* ∈ [0, 1], the forward process evolves according to:

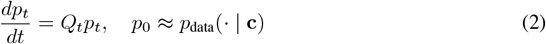

where *Q*_*t*_ ∈ ℝ ^| *χ* | × | *χ* |^ is the token-level diffusion matrix. Each element *Q*(*i, j*) of the matrix *Q* represents the probability that the token at position *i* transitions into the token value at position *j* which defined over the vocabulary. We use an *absorbing* diffusion structure where all tokens transition toward a special ‘MASK’ state. We use a log-linear noise schedule *Q*_*t*_ = *σ*(*t*)*Q*^absorb^ where 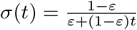 and *ε* = 10^−3^.

### Reverse Process via Concrete Scores

The reverse diffusion process is characterized by the time-reversed matrix 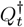 with entries:

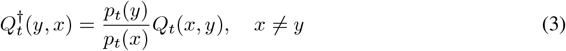

The ratios 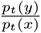 are called *concrete scores* [Meng et al. (2022)], *the discrete analog of* ∇_*x*_ log *p*_*t*_(*x*) in continuous diffusion. For gene expression sequences, we parameterize a score network *s*_*θ*_ : *χ*^*M*^ × [0, 1] →ℝ^*M*×|*χ*|^ that estimates ratios between sequences with Hamming distance 1:

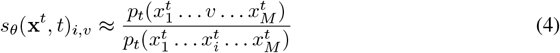

where position *i* has token 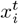 replaced by token *v*. Conditioning on **c** is incorporated by augmenting the network input: *s*_*θ*_(**x**^*t*^, *t*, **c**).

### Training Objective: Denoising Cross-Entropy

Following recent theoretical analysis Ou et al. (2025), for the absorbing case, the DWDSE objective simplifies to a weighted cross-entropy loss. Specifically, the concrete score can be reparameterized as:

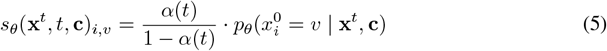

where 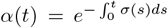 is the survival probability at time *t*. Under this parameterization, the DWDSE loss reduces to:

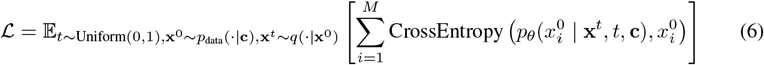

This formulation enables tractable likelihood-based training while naturally handling the discrete, high-dimensional nature of gene expression data.

## 3 Experiments

In this section, we present evaluations that access the effectiveness of proposed DCM-framework to learn and infer cell state representations. To evaluate the quality of generated gene expression profiles, we compare the distribution of predicted gene expression vectors to that of ground-truth expression vectors using two complementary metrics: Maximum Mean Discrepancy (*MMD*) and the 2-Wasserstein distance (*W*_2_). Together, these metrics assess both fine-grained statistical similarity and global geometric alignment between distributions. We use these two metrics to provide a complementary view of model performance. We benchmark our model against SCVI [Lopez et al. (2018a)], scDiffussion [Luo et al. (2024)], CFGen [Palma et al. (2024)], and the current SOTA scDLM [Palla et al. (2025)], with model scores taken from [Palla et al. (2025)].

The **Maximum Mean Discrepancy (***MMD*) is a non-parametric metric that measures the distance between two distributions by comparing their expectations under a rich class of functions. In our setting, *MMD* quantifies how well the distribution of predicted gene expression vectors matches the distribution of true gene expression vectors. Given ground-truth samples *X* = {*x*_1_, *x*_2_, …, *x*_*m*_} and predicted samples *Y* = {*y*_1_, *y*_2_, …, *y*_*m*_}, we compute the unbiased empirical estimator:

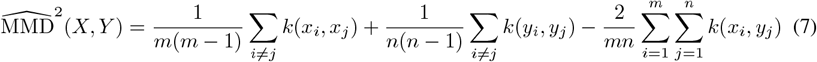

and we use a Gaussian RBF kernel here: 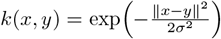 which makes *MMD* sensitive to differences in higher-order statistics and multimodal structure.

The **2-Wasserstein distance** measures the discrepancy between distributions based on optimal transport and directly captures the geometry of the gene expression space. Intuitively, it quantifies the minimum cost of transporting mass from the predicted distribution to the ground-truth distribution. For empirical distributions with equal sample size *n*, this reduces to a discrete matching problem:

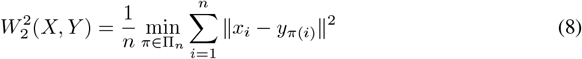

where Π_*n*_ is the set of all permutations of {1, …, *n*}. When both predicted and true gene expression distributions are approximated as Gaussians with means *µ*_*P*_, *µ*_*Q*_ and covariances ∑_*P*_,∑_*Q*_, we use the closed-form expression:

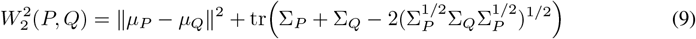

which coincides with the Fréchet Distance (FD). In our evaluation, a lower *W*_2_ indicates that predicted gene expression vectors align closely with ground-truth vectors in terms of global location and covariance structure. We note that some prior works report metrics computed in PCA-reduced space Palma et al. (2024). Since DCM operates directly on raw gene expression counts without dimensionality reduction, we evaluate all metrics in the full gene expression space for consistency.

Our experiments involve two major directions of evaluations: unconditional and conditional generative performances on both observational and perturbational datasets. We employ a DiT backbone Peebles & Xie (2023) for the score network where the diffusion-time condtioning is done via Adaptive LayerNorm (AdaLN). Here we introduce other conditioning variables to the score network as we later describe. We use full gene expression profiles for a cell type as the context length of the transformer (≈17*k*) and employ Flash Attention Dao et al. (2022) for efficient attention computation. To handle variable gene subsets across cells: we use special ‘PAD’ tokens for non-expressed or non-selected genes and mask attention to these tokens to prevent them from influencing the model’s computations. To reduce training time, we use lower precision representations of the gene expression.

### 3.1 Unconditional generation

For this experiment, we are interested in evaluating the model’s fundamental ability to learn and reproduce the underlying distribution of single-cell gene expression data without guidance from conditioning labels with the only conditioning to the score network being the diffusion-time embeddings. Here, we use single cell RNA-sequencing data from the datasets used in both [Palla et al. (2025)], and [Palma et al. (2024)]. We evaluate the model’s output to match the distribution of expression vectors to real cell states that were held out during the training of the model.

#### 3.1.1 results & discussion

DCM achieves substantial improvements over all baselines on both metrics. On *W*2 distance, DCM (5.913) reduces the gap to the ground truth nearly 2-fold compared to scLDM (10.615), the strongest continuous baseline. On *MMD*^2^*RBF*, DCM (0.019) achieves a 5-fold improvement over the closest baseline CFGen (0.075) and a further improvement over scLDM (0.102).

We offer two candidate explanations for these gains, which remain speculative without further ablation. First, single-cell data is zero-inflated, and discrete models represent zeros as a distinct state rather than requiring a sharp mode at zero in continuous space. Second, for lowly-expressed genes, neighboring count values (0, 1, 2) carry categorical significance that continuous interpolation may obscure. Whether DCM’s gains are concentrated in these regimes is an empirical question we leave to future work.

DCM also achieves these results with a 5M parameter model, which is substantially smaller than scLDM’s two-stage architecture (transformer VAE + diffusion transformer). The end-to-end discrete diffusion framework eliminates the need for a separate encoder-decoder pipeline, simplifying training and reducing the number of design choices.

### 3.2 Conditional generation

In this experiment, we train our model to generate gene expression conditioned on multiple attributes: cell type or perturbation type (gene knockouts) or both. For cell types, we create conditioning embedding via one-hot encoding of a specific cell type followed by a linear layer. For perturbation type, we employ a protein language model Lin et al. (2023) to obtain the embeddings for each of the perturbation labels in the dataset, enabling generalisation on held-out labels. The conditioning embeddings are then concatenated with the diffusion-time embeddings and provided to the score network via the AdaLN mechanism. At inference time, DCM is queried with combinations of cell type and perturbations to generate new gene expression profiles.

**Figure 2.**
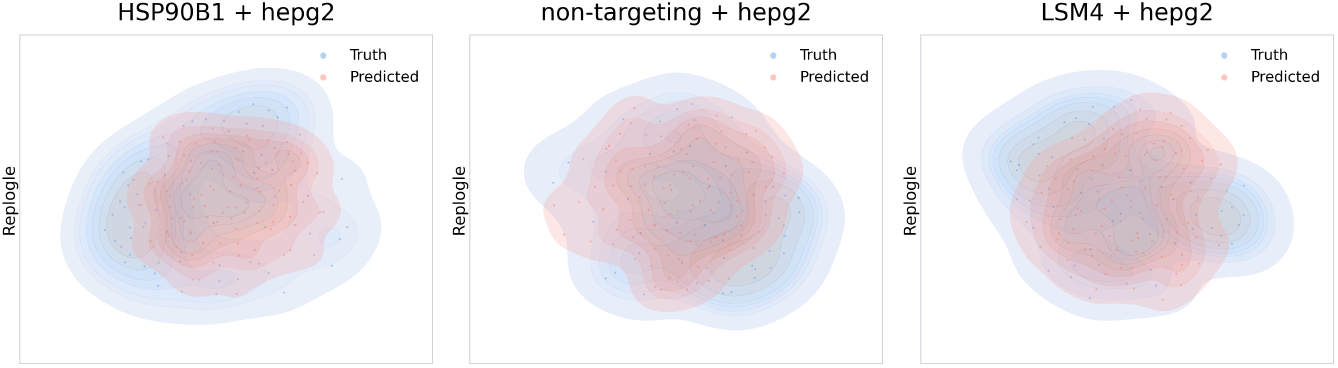
UMAPs from Replogle: (Left) For the HSP90B1 & (Right) LSM4 perturbation, in HepG2 cells, predicted (red) and true (blue) density contours show strong overlap, indicating that the model captures the perturbation-specific transcriptional shift. (Center) Non-targeting control in HepG2 cells confirms the model reproduces the unperturbed baseline distribution.

#### 3.2.1 Results & discussion

In table 2, DCM performs the best on both the *W*_2_ and the *MMD*^2^*RBF* metrics by margins of ≥50% and 7% (5.428 vs 12.457 & 0.020 vs 0.027 for scDLM at *w* = 1) respectively for the Parse 1M benchmark. DCM achieves the best *W*_2_ distance on the full Replogle benchmark (10.03 vs. 11.292 for scLDM at *w* = 1), representing an 13% improvement in global distributional alignment. On the K562-only evaluation, which isolates perturbation modeling from cross-cell-line heterogeneity in the Replogle dataset, DCM achieves a competetive *W*_2_ of 7.284. These results demonstrate that discrete diffusion captures population-level transcriptomic structure effectively in the conditional setting.

**Table 1:**
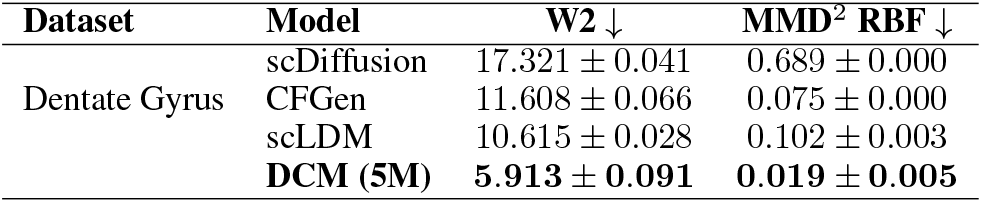
Model generation performance on unconditional cell generation benchmarks, for the dentate gyrus dataset.

**Table 2:** Model generation performance on conditional perturbation prediction benchmarks (Replogle dataset). DCM is evaluated at two scales: 20M parameters on the full dataset (4 cell lines) and 10M parameters on K562 only. The K562-only setting isolates perturbation modeling from cross-cell-line variation.

| Dataset | Model | $W_2 \downarrow$ | $MMD^2 \text{ RBF} \downarrow$ |
| --- | --- | --- | --- |
| Parse 1M | scVI | $35.508 \pm 0.182$ | $1.372 \pm 0.016$ |
| | CPA | $13.534 \pm 0.036$ | $1.117 \pm 0.014$ |
| | scGPT | $22.870 \pm 0.152$ | $2.203 \pm 0.013$ |
| | STATE | $19.111 \pm 0.136$ | $0.714 \pm 0.009$ |
| | scLDM ($w = 1$) | $12.457 \pm 0.045$ | $0.027 \pm 0.002$ |
| | scLDM ($w = 5$) | $12.902 \pm 0.087$ | $0.071 \pm 0.004$ |
| | scLDM ($w = 10$) | $13.638 \pm 0.111$ | $0.122 \pm 0.006$ |
|  | **DCM (10M)** | **$5.428 \pm 0.163$** | **$0.020 \pm 0.012$** |
| Replogle | scVI | $17.359 \pm 0.051$ | $0.453 \pm 0.003$ |
| | CPA | $11.510 \pm 0.029$ | $0.532 \pm 0.003$ |
| | scGPT | $34.166 \pm 0.272$ | $3.087 \pm 0.010$ |
| | STATE | $20.58 \pm 0.039$ | $0.730 \pm 0.003$ |
| | scLDM ($w = 1$) | $11.292 \pm 0.033$ | **$0.200 \pm 0.002$** |
| | scLDM ($w = 5$) | $12.900 \pm 0.069$ | $0.320 \pm 0.004$ |
| | scLDM ($w = 10$) | $14.911 \pm 0.091$ | $0.436 \pm 0.005$ |
| | **DCM (20M)** | **$10.03 \pm 0.337$** | $0.688 \pm 0.099$ |
| Replogle - K562 | **DCM (10M)** | $7.284 \pm 0.091$ | $0.605 \pm 0.005$ |

However, we observe a divergence between *W*_2_ and *MMD*^2^*RBF* on these benchmarks: DCM’s *MMD*^2^ (0.688) is higher than scLDM’s (0.200) for Replogle and is comparable for the Parse 1M benchmarks. The *W*_2_ metric (Eq. 9) captures mean and covariance alignment—the first two moments of the distribution. The *MMD*^2^*RBF* metric is sensitive to higher-order distributional features, including gene-gene correlations and tail behavior. DCM’s strong *W*_2_ but weaker *MMD*^2^*RBF* therefore suggests that it accurately recovers the mean expression profile and gene-level variances for each perturbation condition, but introduces errors in higher-order dependency structure.

We consider three candidate explanations for this pattern. First, our conditioning mechanism, additive embedding concatenation, cannot represent interactions between perturbation and cell type. The same knockdown produces different effects in different cell lines, and capturing this requires multiplicative or attention-based conditioning; scLDM’s cross-attention mechanism may provide this capacity. Supporting this hypothesis, *MMD*^2^*RBF* improves in the K562-only setting where cell-type interactions are absent, and is strong on unconditional Dentate Gyrus where no conditioning is required. Second, gene-gene correlations induced by perturbations may be inherently better captured in continuous latent spaces, which provide smooth interpolation between related states. The *MMD*^2^*RBF* gap persists even on K562 alone (0.605 vs. scLDM’s 0.200), though conditioning differences confound this comparison. Third, the SEDD training objective may be less effective than flow matching for capturing correlational structure, independent of the discrete-versus-continuous distinction.

Overall, discrete diffusion achieves state-of-the-art performance on *W*_2_ for conditional perturbation prediction, demonstrating that the approach extends effectively from unconditional to conditional generation. Ablations to isolate the source of the *MMD*^2^*RBF* gap are left to future work. We also note that the sequencing-depth of the datasets need to be comparable for discrete modelling.

## 4 Conclusion

In this work, we demonstrated that enforcing the discreteness of raw transcriptomic data in diffusion frameworks improves generative modeling in unconditioned and conditioned settings. We introduced DCM, a discrete diffusion model that achieves new state-of-the-art performance in unconditioned generation on the Dentate Gyrus dataset, indicating the model has formed better representations of global and granular transcriptomic patterns. On the Replogle dataset, DCM similarly improves *W*_2_, confirming stronger cell population-level alignment, while performance on *MMD*^2^*RBF* is more variable. This suggests that while discrete diffusion robustly captures global transcriptomic structure across datasets, fine-grained dependency modeling may depend on conditioning complexity, highlighting an important direction for future work.

Beyond the empirical improvements, this work demonstrates that generative models gain representational power when their state space matches the discrete, sparse structure of the biological measurements they aim to model. While applied here with single-cell transcriptomics data, this principle also extends to other count-based molecular assays that will enable more faithful virtual cells.

## A Appendix: DATASETS USED

**Table 3:**
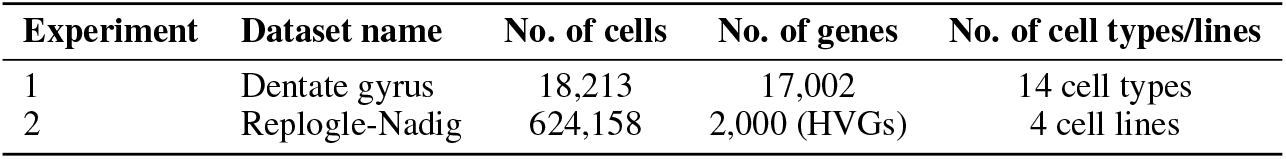
Summary of datasets used in the experiments.

## B Appendix: Training details

**Table 4:**
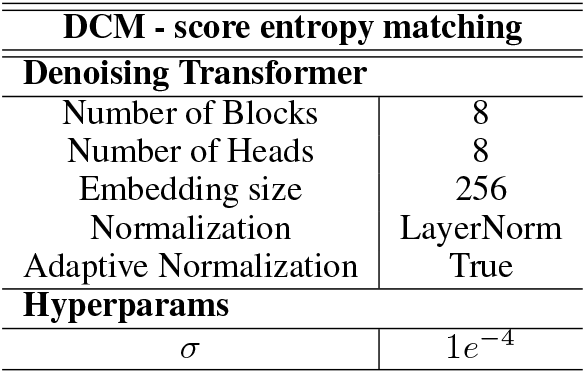
Hyperparameter values of DCMs considered in this paper.

## C CODE

The implementation of the methods discussed in this paper is available online at [https://github.com/sanjukta7/aivc-dcm].

